# Synaptic and peptidergic connectome of a neurosecretory centre in the annelid brain

**DOI:** 10.1101/115204

**Authors:** Elizabeth A. Williams, Csaba Verasztó, Sanja Jasek, Markus Conzelmann, Réza Shahidi, Philipp Bauknecht, Gáspár Jékely

## Abstract

Neurosecretory centres in animal brains use peptidergic signalling to influence physiology and behaviour. Understanding neurosecretory centre function requires mapping cell types, synapses, and peptidergic networks. Here we use electron microscopy and gene expression mapping to analyse the synaptic and peptidergic connectome of an entire neurosecretory centre. We mapped 78 neurosecretory neurons in the brain of larval *Platynereis dumerilii*, a marine annelid. These neurons form an anterior neurosecretory organ expressing many neuropeptides, including hypothalamic peptide orthologues and their receptors. Analysis of peptide-receptor pairs revealed sparsely connected networks linking specific neuronal subsets. We experimentally analysed one peptide-receptor pair and found that a neuropeptide can couple neurosecretory and synaptic brain signalling. Our study uncovered extensive non-synaptic signalling within a neurosecretory centre and its connection to the synaptic brain.

## Introduction

Nervous system signalling occurs either at synapses or via secreted diffusible chemicals that signal to target cells expressing specific receptors. Synapse-level connectomics using electron microscopy allows mapping synaptic networks, but fails to reveal non-synaptic signalling. In addition to acting in a neuroendocrine fashion, non-synaptic volume transmission by neuropeptides and monoamines can have neuromodulatory effects on synaptic signalling (Bargmann 2012; Marder 2012). Overlaying synaptic and peptidergic maps is challenging and requires knowledge of the expression of the modulators and their specific receptors as well as synaptic connections. Such mapping has only been achieved for relatively simple circuits, such as the stomatogastric nervous system of crustaceans where synaptic connections are known and the effect of neuropeptides and the activation of single peptidergic neurons can be analysed experimentally (Stein et al. 2007; Thirumalai and Marder 2002; Blitz et al. 1999). Likewise, connectome reconstructions combined with cellular-resolution neuropeptide and receptor mapping allow the dissection of peptidergic signalling in *Drosophila* (Schlegel et al. 2016). In *Caenorhabditis elegans,* the spatial mapping of monoamines and neuropeptides and their G protein-coupled receptors (GPCRs) revealed interconnected networks of synaptic and non-synaptic signalling (Bentley et al. 2016).

Neurosecretory centres, such as the vertebrate hypothalamus and the insect ring gland, found in the anterior brain of many animals show exceptionally high levels of neuropeptide expression (Herget and Ryu 2015; Siegmund and Korge 2001; Campbell et al. 2017) suggesting extensive non-synaptic signalling. These centres coordinate many processes in physiology, behaviour, and development, including growth, feeding, and reproduction (Sakurai et al. 1998; Bluet-Pajot et al. 2001; Sternson et al. 2013). The combined analysis of synaptic and peptidergic networks is particularly challenging using single marker approaches for neurosecretory centres that express dozens of neuropeptides. To fully understand their function, a global mapping of peptidergic networks within these brain regions and how they connect to the rest of the nervous system is required.

Here, we analyse synaptic and peptidergic signalling in the anterior neurosecretory centre in larval *Platynereis dumerilii,* a marine annelid. Annelid and other marine larvae have an anterior sensory centre, the apical organ, involved in the detection of various environmental cues (Hadfield et al. 2000; Conzelmann et al. 2013; Page 2002; Chia and Koss 1984). The apical organ is neurosecretory and expresses diverse neuropeptides that are thought to regulate various aspects of larval behaviour and physiology, including the induction of larval settlement and metamorphosis (Mayorova et al. 2016; Thorndyke et al. 1992; Tessmar-Raible et al. 2007; Conzelmann et al. 2011; Marlow et al. 2014). Apical organs have a conserved molecular fingerprint across marine larvae, suggesting that they represent a conserved sensory-neuroendocrine structure (Marlow et al. 2014). The apical organ area or apical nervous system (ANS) in *Platynereis* larvae shows molecular similarities to other neuroendocrine centres, including the pars intercerebralis in insects and the vertebrate hypothalamus, suggesting a common ancestry (Tessmar-Raible et al. 2007; Steinmetz et al. 2010; Conzelmann et al. 2013). Molecular and developmental similarities in various protostomes and deuterostomes further suggest a more widespread conservation of neuroendocrine centres (Hartenstein 2006; Wirmer et al. 2012; Tessmar-Raible 2007). The study of marine invertebrate larval apical organs could thus inform about the evolution of neuroendocrine cell types and signalling mechanisms in metazoans.

*Platynereis* larvae represent a powerful system to analyse gene expression and synaptic connectivity in a whole-body context, allowing linking distinct neuropeptides and other molecules to single neurons (Asadulina et al. 2012; Williams and Jékely 2016;Shahidi et al. 2015; Achim et al. 2015; Pettit et al. 2014; Vergara et al. 2017). To understand how synaptic and peptidergic signalling is integrated in the *Platynereis* ANS, we combine serial section electron microscopy with the cellular analysis of neuropeptide signalling. This combined analysis revealed extensive non-synaptic peptidergic signalling networks within the ANS distinguishing this area from the rest of the nervous system. Through connectomics and functional studies we also reveal how this endocrine region can interact with the synaptic nervous system by peptidergic modulation of the ciliomotor circuitry.

## Results

### Ultrastructural reconstruction of the anterior neurosecretory centre

To comprehensively map a neurosecretory area with ultrastructural detail, we focused on the larvae of *Platynereis.* Due to their small size, the larvae are amenable to whole-body connectomic analysis (Randel et al. 2015; Shahidi et al. 2015). We used a full-body serial electron microscopy dataset of a 3-day-old larva (Randel et al. 2015) and reconstructed its entire apical neurosecretory nervous system (Figure 1A-D). *Platynereis* and other annelids have an anterior neurosecretory plexus containing the projections of peptidergic sensory-neurosecretory neurons (Tessmar-Raible et al. 2007; Aros et al. 1977). The neurosecretory plexus forms an anatomically and ultrastructurally distinct area that can be clearly distinguished from other neuropils, including the adjacent optic and nuchal organ neuropils (Randel et al. 2014; Shahidi et al. 2015) (Figure 1D). Neurites in this area have a high number of dense-core vesicles and very few synapses (Figure 1E, F). Classic neurotransmitter synapses can be identified by large clusters of clear synaptic vesicles in the axons, while peptidergic synapses appear as smaller clusters of dense core vesicles (Randel et al. 2014; Shahidi et al. 2015). We reconstructed all neurons that project to this region (Figure 1B, Supplement 1 to figure 1, Video 1) and identified 70 sensory neurons and eight projection interneurons, the latter project in and out of the neurosecretory plexus. Most of the sensory neurons are bilaterally symmetric pairs with distinct axonal projection patterns, except for a few asymmetric neurons (Supplement 1 to figure 1). The sensory neurons have diverse apical sensory specializations. Based on these morphological criteria we could distinguish at least 20 different sensory cell types with likely different sensory functions. For example, there are four ciliary photoreceptor cells (cPRC)(Arendt et al. 2004) with highly extended ciliary membranes, one asymmetric neuron with five sensory cilia (SN^YFa5cil^), a pair of neurons with two long parallel cilia (SN^WLD1^), a pair with long branched sensory cilia (SN^NS29^), and three uniciliary neurons that are part of the nuchal organ (SN^nuchNS^), a putatively chemosensory annelid organ (Purschke 1997). Twenty-five uniciliated neurons (23 SN^DLSO^ cells, SN^PDF^-^dcl2^, and SN^PDF-dcr3^) are part of a dorsolateral sensory cluster (Supplement 1 to figure 1). Most of the sensory neurons have axonal projections that are extensively branched within the neurosecretory plexus (Supplement 1 to figure 1) and that are filled with dense-core vesicles (Figure 1F). The pairs of left-right symmetric sensory neurons project to similar areas of the neurosecretory plexus revealing a fine-scale organization within the plexus (Figure 1G and Supplement 1 to figure 1, Video 1). We refer to all neurons that project to the neurosecretory plexus as the apical nervous system (ANS).

**Figure 1.**
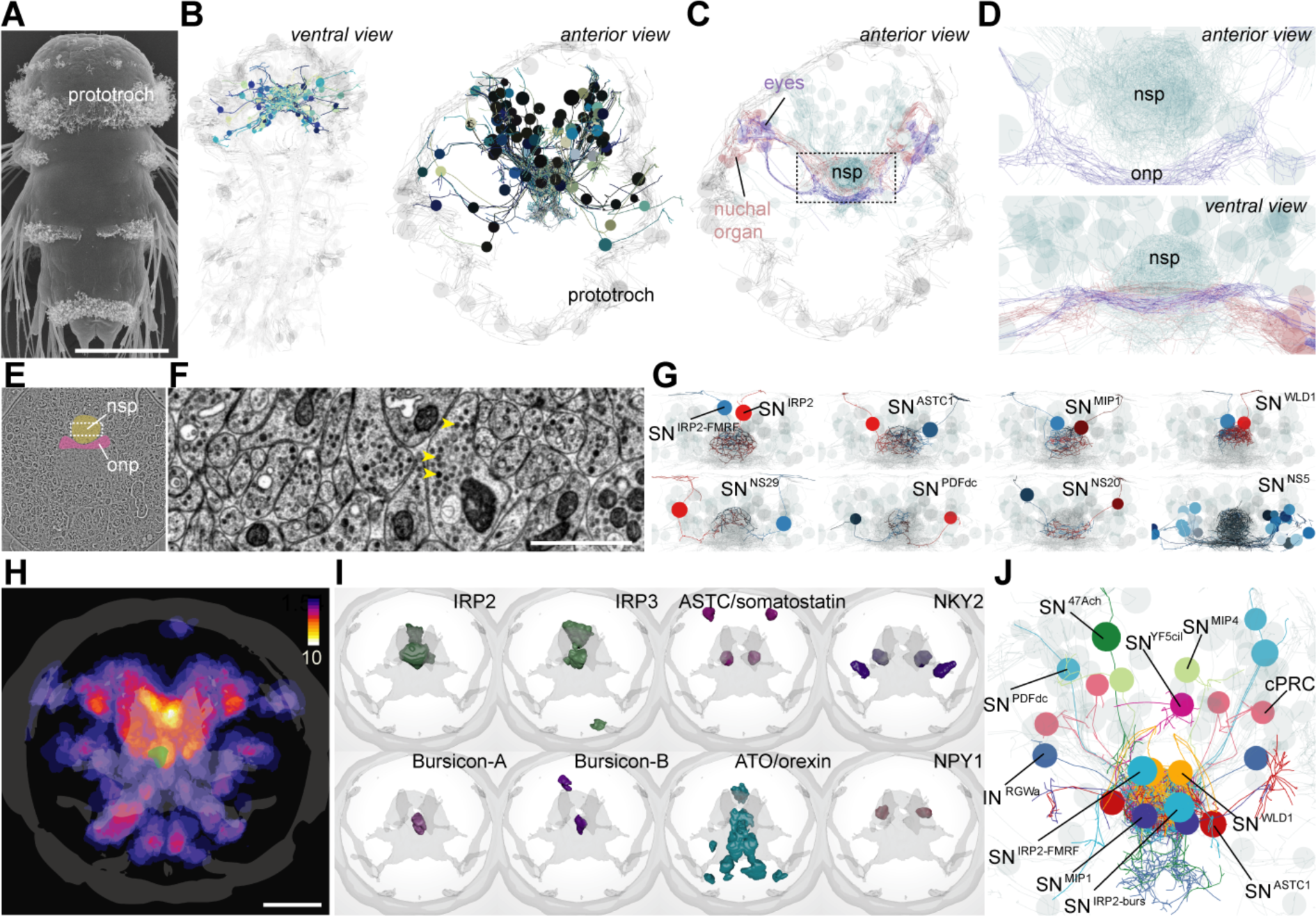
EM reconstruction and mapping of proneuropeptide expression in the apical nervous system of *Platynereis* larvae. (**A**) Scanning electron micrograph of a 3-day-old *Platynereis* larva. (**B**) Reconstructed apical nervous system neurons (ANS) (shades of blue) in a full body transmission electron microscopy (TEM) dataset of a 3-day-old larva, shown against a framework of reconstructed ciliated cells and axonal scaffold (light grey). Axons and dendrites appear as lines and cell body positions are represented by spheres. (**C**) ANS neurons project to an apical neurosecretory plexus in the centre of the head, which forms a small sphere dorsal and apical to the optic neuropil and nuchal organ neuropil. (**D**) Close-up view of neurosecretory plexus indicated by box in C. (**E**) TEM image of a section in the larval head. The neurosecretory plexus and the optic neuropil are highlighted. Boxed area is shown in F. (**F**) Close-up view of the neurosecretory plexus. Yellow arrowheads point at dense core vesicles. (**G**) Examples of bilaterally symmetric pairs of sensory neurons that innervate different regions of the neurosecretory plexus in the reconstructed ANS (grey), ventral view. (**H**) Heat map of the expression of 48 proneuropeptides in the 2- day-old *Platynereis* larva, apical view, projected on a reference scaffold of cilia and axonal scaffold (grey). Colour scheme indicates number of different proneuropeptides expressed in a cell. An apical FMRFamide-expressing neuron of the ANS (SN^IRP2-FMRF^) is overlaid for spatial reference (green). (**I**) Examples of average gene expression patterns of individual proneuropeptides from the 2-day proneuropeptide expression atlas. (**J)** TEM reconstruction of the 3-day-old larval ANS showing neurons that could be assigned specific proneuropeptide expression based on position and sensory cilia morphology. Abbreviations: nsp, apical neurosecretory plexus; onp, optic neuropil. Scale bars: A, 70 μm; H, 30 μm; F, 1 μm.

### Comprehensive mapping of neuropeptide expression in *Platynereis* larvae

The *Platynereis* larval ANS is known to express many neuropeptides, including vasotocin, FMRFamide, and myoinhibitory peptide (Tessmar-Raible et al. 2007; Conzelmann et al. 2011; Conzelmann et al. 2013). To comprehensively analyse neuropeptide expression in the whole larva, we used whole-mount *in situ* hybridization for 51 *Platynereis* proneuropeptides (of 98 total (M Conzelmann et al. 2013)). We used image registration to spatially map all neuropeptide expressions to a common nuclear reference template (Asadulina et al. 2012). We summed all binarized average expression domains and found that the region with the highest proneuropeptide expression corresponds to the ANS (Figure 1H, Figure 1 – source data 1). Some voxels in the map coexpress up to 10 different neuropeptides. Neuropeptides that were expressed in the ANS include two *Platynereis* insulin-like peptides (IRP2, IRP3), two bursicons (bursicon-A, -B), achatin, myoinhibitory peptide (MIP), and several homologs of hypothalamic peptides (Mirabeau and Joly 2013; Jékely 2013) including NPY (three homologs, NPY1, NPY4, NKY2), orexin/allatotropin, tachykinin, galanin/allatostatin-A, and allatostatin-C/somatostatin (Figure 1I and Supplement 2 to figure 1).

The acetylated tubulin antibody we used to counterstain the *in situ* samples labels cilia and axonal scaffold. With this counterstaining signal, we correlated neurons expressing specific proneuropeptides and with distinct ciliation to sensory ANS neurons reconstructed from EM data (Figure 1J and Supplement 3 to figure 1). For other ANS neurons (SN^PDFdc^, IN^RGW^) we assigned neuropeptides based on direct immunogold labelling on the same EM series (Shahidi et al. 2015). Overall, we mapped neuropeptide expression to 25 reconstructed ANS neurons, including sensory neurons coexpressing IRP2 and FMRFamide (SN^IRP2-FMRF^), IRP2 and bursicon (SN^IRP2- burs^), or expressing ASTC/somatostatin (SN^ASTC1^) (Figure 1J and Supplement 3 to Figure).

### Low level of synaptic connectivity within the ANS

We next analysed how the ANS neurons are synaptically connected. We found that most neurons have no or only very few synapses (Figure 2, Supplementary table 1). 33% of the neurons have 0-2 synapses despite highly branched axonal projections filled with dense core vesicles (Figure 2). This suggests that these neurons predominantly use volume transmission. The synapses we identified in most neurons contained dense-core vesicles, indicative of their peptidergic nature. The four cPRCs, an asymmetric sensory neuron (SN^47Ach^), and the 8 projection neurons have the highest number of synapses. In addition, 15 ANS sensory neurons have 10 or more peptidergic synapses. The cPRCs express a cholinergic marker (Jékely et al. 2008) and contain large synapses with clear vesicles. The other sensory cells have ultrastructurally distinct, small peptidergic synapses characterized by dense core vesicles clustering at the membrane (Figure 2).

**Figure 2.**
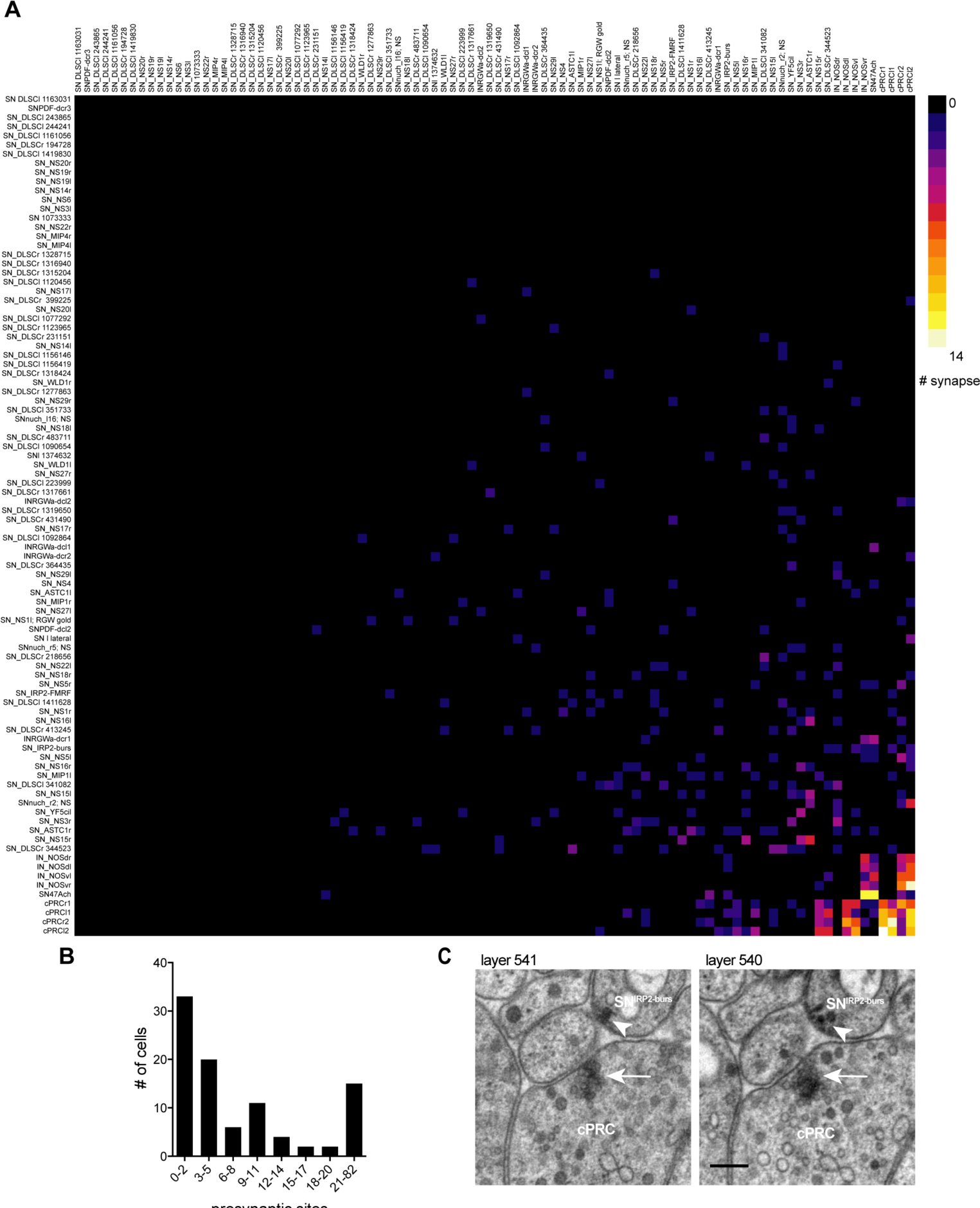
Low level of synaptic connectivity of ANS neurons. (**A**) Full matrix of synaptic connectivity of ANS neurons. With the exception of the ciliary photoreceptors, a few sensory cells, and the IN^NOS^ interneurons, the ANS shows very sparse synaptic connectivity. (**B**) Histogram of the number of presynaptic sites in ANS neurons. (**C**) Comparison of a cholinergic synapse of a ciliary photoreceptor (arrow) to a peptidergic synapse of the SN^IRP2-burs^ cell (arrowhead). Two consecutive sections are shown. Scale bar in (**C**), 200 nm.

### Analysis of peptidergic signalling networks in the ANS

The high number of neuropeptides expressed in the ANS and the low degree of synaptic connectivity of the cells prompted us to further analyse peptidergic signalling networks in the *Platynereis* larva. We used two resources: an experimentally determined list of *Platynereis* neuropeptide GPCRs (Bauknecht and Jékely 2015) and a spatially-mapped single-cell transcriptome dataset (Achim et al. 2015). The GPCR list included 18 deorphanized neuropeptide receptors, to which we added a further three deorphanized neuropeptide receptors. We deorphanized a GnRH receptor activated by both *Platynereis* GnRH1 and GnRH2 (M Conzelmann et al. 2013), and second receptors for vasotocin and myomodulin (Supplement 1 to figure 3). The single-cell transcriptome data consisted of cells of the head (episphere) of 2-day-old larvae. The cells were mostly neurons (Achim et al. 2015).

To comprehensively analyse potential peptide-receptor signalling networks, we first created a virtual larval brain with cells arranged in an approximate spatial map (Figure 3A, Supplement 2 to figure 3, Supplementary Table 1, Figure 3 – source data 1, Figure 3 – source data 2). To this virtual map, we mapped the expression of all proneuropeptides and deorphanized receptors (Figure 3C, D). The combined expression of 80 proneuropeptides showed a similar pattern to the *in situ* map with a highly peptidergic group of cells in the ANS region, as defined by the endocrine marker genes *Phc2, dimmed, Otp* and *nk2.1* (Conzelmann et al. 2013; Tessmar-Raible et al. 2007) (Figure 3B). Given the high level of peptide expression in the ANS, the mapping of peptidergic signalling networks will mostly reveal potential signalling partners within this region or from the ANS to the rest of the brain. Most GPCR expression was also concentrated in the ANS in peptidergic cells. Several cells expressed a unique combination of up to 9 GPCRs (Figure 3D).

**Figure 3.**
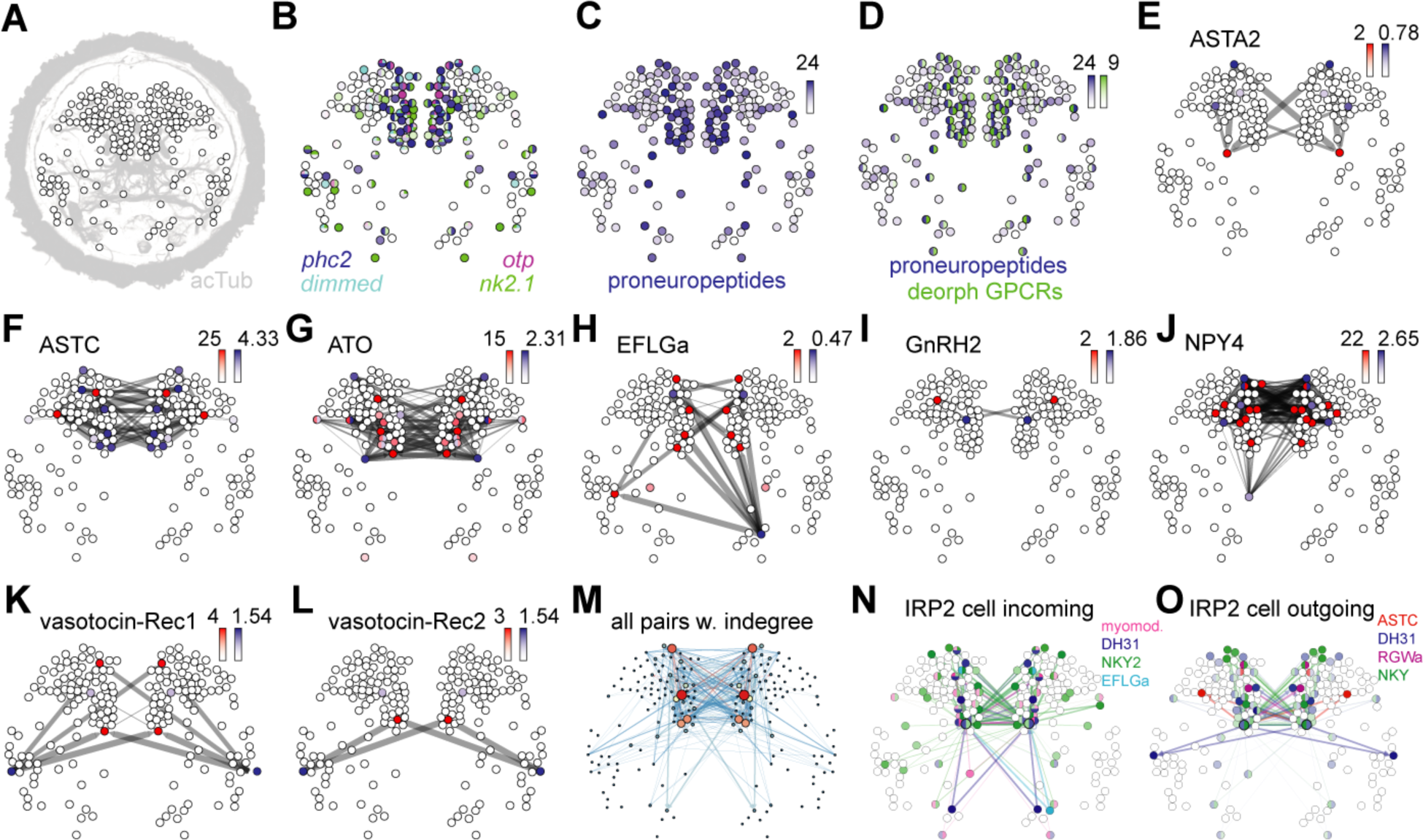
Mapping the peptidergic connectome in the *Platynereis* larval head. (**A**) Positions of cells (nodes) from single cell RNA-Seq from 2-day-old larvae placed in an approximate spatial map, projected on an acetylated tubulin immunostaining (grey, anterior view). Samples with predicted bilateral symmetry were represented as two mirror image nodes in the map. (**B**) Expression of neuroendocrine marker genes projected on the single cell map. Colour intensity of nodes reflects magnitude of normalized log_10_ gene expression. (**C**) Map of combined expression of 80 proneuropeptides expressed in each single cell sample. (**D**) Map of combined expression of 80 proneuropeptides and 23 deorphanized GPCRs expressed in each single cell sample. (**E–L**) Connectivity maps of individual neuropeptide-GPCR pairs, coloured by weighted in-degree (red) and proneuropeptide log_10_ normalized expression (blue). Arrows indicate direction of signalling. Arrow thickness determined by geometric mean of log_10_ normalized proneuropeptide expression of signalling cell and log_10_ normalized GPCR expression of corresponding receiving cell. (**M**) Connectivity map of all possible known neuropeptide-GPCR signalling, colour and node size represent weighted in-degree. (**N**) Connectivity map of an IRP2-expressing cell with incoming neuropeptide signals. (**O**) Connectivity map of an IRP2-expressing cell with outgoing neuropeptide signals.

We also used the spatially mapped single cell transcriptome data to map the expression of different types of sensory receptor genes in the opsin and transient receptor potential (Trp) channel families. These receptor genes are known for their function in the detection and transduction of light/pain/temperature/mechanical stimuli (Terakita 2005; Moran et al. 2004). Neurons in the ANS of 2-day-old larvae express diverse combinations of these genes (Supplement 3 to figure 3). This supports our conclusion from the morphological data that these neurons are responsible for detecting a variety of different environmental cues.

Coexpression analysis with small neurotransmitter synthesis enzymes revealed 1 cell (1% of total) with only neurotransmitter markers but no neuropeptide, 76 (71% of total) purely peptidergic cells, and 21 cells that coexpress small transmitters and neuropeptides (19.6% of total)(Supplement 4 to figure 3). The remaining 9 cells expressed neither neuropeptides nor neurotransmitter markers and thus are likely non-neuronal cells (8.4% of total). Strikingly, 2 ANS cells coexpress up to 24 different proneuropeptides. Based on specific neuropeptide expression and the spatial mapping we could correlate 15 cells to ANS cells reconstructed from EM data (Supplementary Table 1). This was possible despite the two resources being derived from different larval stages, because many ANS cells are already differentiated in 2-day-old larvae and readily identifiable between stages.

To establish peptidergic signalling networks, we treated peptide-expressing cells as source nodes and GPCR-expressing cells as target nodes. We define edge weights in the directed graphs as the geometric mean of normalized proneuropeptide expression in the source and normalized GPCR expression in the target. This way, we also consider the expression level of peptides and receptors. Receptor expression can correlate with neurophysiological sensitivity to a neuropeptide (Garcia et al. 2015; Root et al. 2011). For each peptide-receptor pair, we projected these networks onto the virtual map (Figure 3E–L, Supplement 4 to figure 3, Figure 3 - source data 3). The chemical connectivity maps of individual ligand-receptor pairs show very sparse and specific chemical wiring. On average, less than 1% of all potential connections are realized (0.8% graph density averaged for all peptide-receptor pairs). Single cells expressing many different neuropeptides generally link to non-overlapping target nodes by each peptide-receptor channel (Figure 3O and Supplement 4 to figure 3Q). Conversely, multiple signals can converge on one cell that expresses more than one GPCR (Figure 3N).

The combined multichannel peptidergic connectome of 23 receptor-ligand pairs forms a single network with an average clustering coefficient of 0.49 and an average minimum path length of 1.54, forming a small-world network (Watts and Strogatz 1998). Analysis of the combined network revealed highly connected components, including nodes that can act as both source and target as potential mediators of peptide cascades (Figure 3N, O). The three neuron types with the highest weighted in-degree and authority value include a pair of dorsal sensory neurons, a central pair of IRP2 neurons, and a pair of RGWamide-expressing neurons (Figure 3M and Figure 4A). The IRP2 neurons are under the influence of an NPY peptide and EFLGamide, the *Platynereis* homolog of thyrotropin-releasing hormone (Bauknecht and Jékely 2015) and express 20 different proneuropeptides.

**Figure 4.**
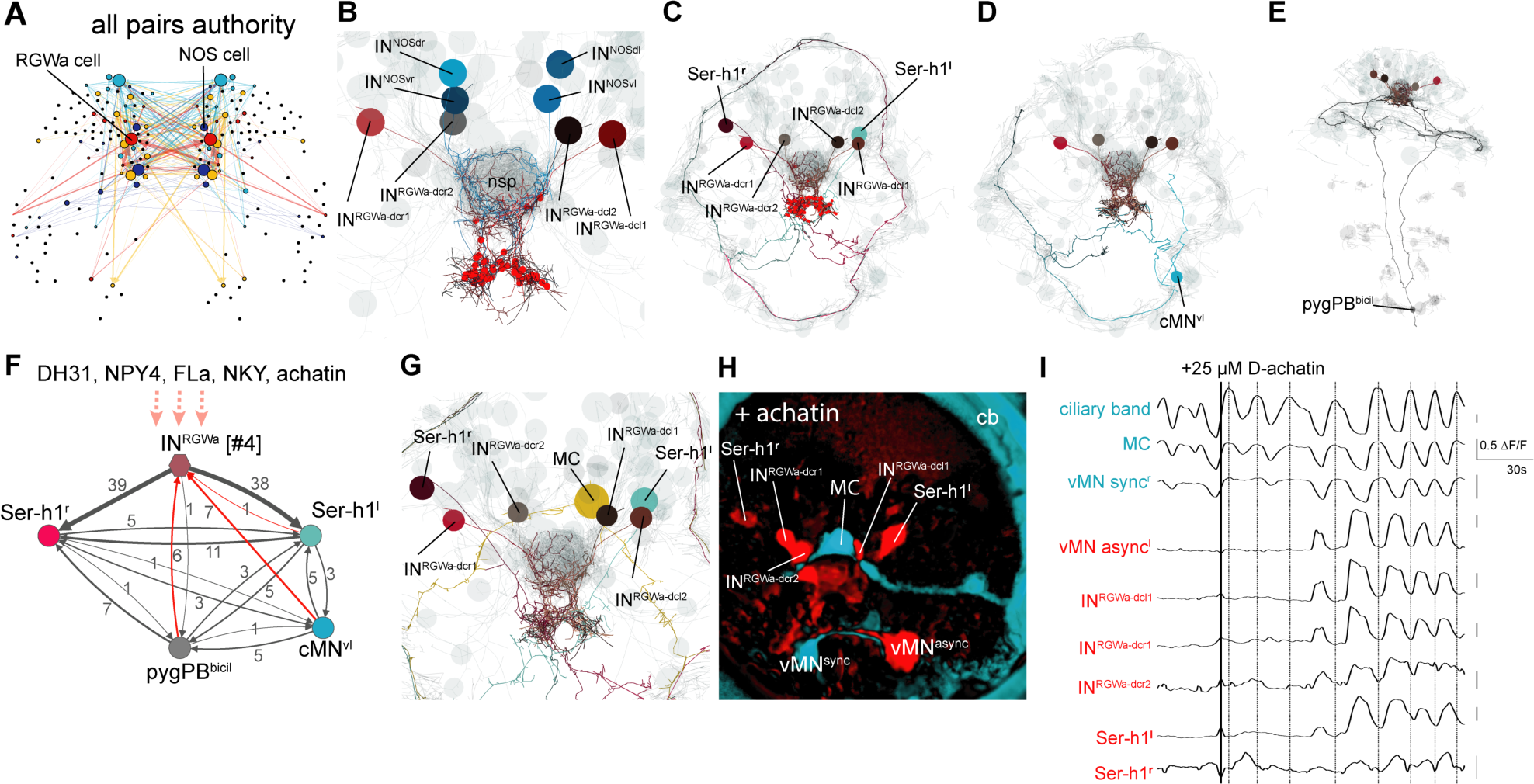
Projection neurons connect the ANS to the synaptic nervous system. (**A**) Connectivity map of all possible known neuropeptide-GPCR signalling, colours represent modules defined by randomized community detection analysis in Gephi and node size represents authority. (**B**) Reconstructed IN^RGWa^ projection neurons (ANS) (shades of brown) in a full body transmission electron microscopy (TEM) dataset of a 3-day-old larva, shown against a framework of reconstructed ANS neurons (light grey). Axons and dendrites appear as lines and cell body positions are represented by spheres. (**C**) TEM reconstruction of IN^RGWa^ projection neurons and Ser-h1 serotonergic neurons. Red spheres indicated IN^RGWa^ presynaptic sites. (**D**) TEM reconstruction of IN^RGWa^ projection neurons and a presynaptic ciliomotor neuron, cMN^vl^. (**E**) TEM reconstruction of IN^RGWa^ neurons and a presynaptic serotonergic sensory neuron, pygPD^bicil^. (**F**) Synaptic connectivity graph of IN^RGWa^, Ser-h1, pygPD^bicil^, and cMN^vl^ neurons. Synaptic inputs from the synaptic nervous system to the IN^RGWa^ neuron are in red. Peptidergic inputs to the IN^RGWa^ neurons are indicated with dashed arrows. (**G**) TEM reconstruction of IN^RGWa^ projection neurons, Ser-h1 neurons, and the cholinergic ciliomotor MC neuron, anterior view (**H**) Confocal microscopy image of correlated pixels of GcAMP6s signal in a 2-day-old *Platynereis* larva after the addition of 25 μM D-achatin neuropeptide, anterior view. Cells showing correlated activity with the serotonergic neurons (red) and the MC cell (cyan) are shown. IN^RGWa-dcl2^ could not be identified in this larva and is likely obscured by the MC cell and/or IN^RGWa-dcl1^. (**I**) Neuronal activity patterns of individually identified neurons in a 2-day-old larva treated with 25 μM achatin.

### Projection interneurons connect the peptidergic ANS to the synaptic nervous system

The RGWamide-expressing neurons express two NPY receptors (NPY4 and NKY) and an achatin receptor, and 17 proneuropeptides, but no markers for small neurotransmitters. By position and RGWamide-expression we identified these cells as IN^RGW^ projection neurons. We also identified cells that likely correspond to IN^NOS^ projection neurons, based on RYamide expression and expression of *nitric oxide synthase* (unpublished results). The four IN^RGW^ projection interneurons as well as the four IN^NOS^ projection interneurons have a distinct anatomy (Figure 4B). They lack dendrites or cilia and their axons are extensively branched in the neurosecretory plexus and project out of this area to a ventral neuropil. Here the IN^RGW^ cells form many peptidergic synapses on two serotonergic ciliomotor neurons (Ser-h1)(Figure 4C) that are activate during periods of ciliary beating (Verasztó et al. 2017). The IN^NOS^ cells only synapse on the IN^RGW^ cells in the ventral neuropil. The IN^RGW^ neurons also receive synapses in the ventral neuropil area from two non-ANS neurons. These neurons (cMN^vl^ and the serotonergic pygPB^bicil^) belong to the ciliomotor circuit of the larva (Figure 3D-F). From the ANS, the IN^RGW^ projection neurons receive small transmitter synapses from the cPRC cells, mixed clear and dense core vesicle synapses from the SN^47Ach^ neuron in the ventral neuropil, and peptidergic synapses from three other ANS sensory neurons in the neurosecretory plexus (Figure 4, Supplementary Table 1). The distinct anatomy and connectivity as well as the high authority values of the projection neurons in the peptidergic network indicate that these cells are important in relaying peptidergic signals from the ANS to the rest of the brain.

To test how these neurons respond to neuropeptides, we used calcium-imaging experiments. The IN^RGW^ and IN^NOS^ neurons express 7 deorphanized GPCRs (Figure 4F), including a receptor for achatin that is specifically activated by achatin peptide containing a D-amino acid (Bauknecht and Jékely 2015). When we treated larvae with D-achatin, four neurons in the ANS were rhythmically activated. These neurons correspond by position to the IN^RGW^ cells (Figure 4I). The rhythmic activation was in-phase with the activation of the Ser-h1 neurons and a ventral motorneuron (vMN^async^), but out of phase with the main cholinergic motorneuron of the head ciliary band, the MC neuron (readily identifiable by calcium imaging (Verasztó et al. 2017)). We could not observe a similar effect upon L-achatin treatment (data not shown). *Platynereis* larvae have a cholinergic and a serotonergic circuit that oscillate out-of-phase. The cholinergic phase arrests the cilia and the serotonergic phase correlates with ciliary beating (Verasztó et al. 2017). Correlation analysis of neuronal activity patterns revealed that D-achatin coupled the activity of the IN^RGW^ cells to the serotonergic cells, and increased the negative correlation between the activity patterns of serotonergic neurons and the MC neuron (Figure 4I and Supplement 1 to Figure 4). This provides an example of how peptidergic signalling in the ANS could recruit neurons to a rhythmically active circuit, enhance the rhythm, and thereby potentially influence locomotor activity.

## Discussion

Here we presented a comprehensive anatomical description of the ANS in the *Platynereis* larva. We combined this with the cellular-resolution mapping of neuropeptide signalling components to analyse potential peptidergic signalling networks. The use of scRNA-seq has a great potential to reveal such signalling networks but also has limitations. For example, we could only score proneuropeptide and receptor mRNA expression and not protein expression levels, peptide release, or the degree of neuronal activation. We also only analysed a relatively small single-cell dataset derived from the head of the larva and thus could not investigate long-range neuroendocrine signalling from the larval episphere to the rest of the body. The first analyses we present here can be extended when more data become available and can also be applied to other species (e.g., (Campbell et al. 2017)). Nevertheless, our approach can reveal all potential peptidergic connections and allows the development of hypotheses on peptidergic signalling that can be experimentally tested, as we have shown here for achatin.

We found that the *Platynereis* larval ANS has a low degree of synaptic connectivity, a strong neurosecretory character, and shows a high diversity of neuropeptide expression. This suggests that the ANS is wired primarily chemically and not synaptically. We identified very specific peptidergic links that connect only a small subset of ANS neurons. Strikingly, it is not only the proneuropeptides that are expressed with high diversity in the ANS, but also their receptors, suggesting that most peptidergic signalling in the brain occurs within the ANS. A high diversity of neuropeptide receptor expression has also been reported for the mouse hypothalamus (Campbell et al. 2017). These observations suggest that there is more extensive peptidergic integration within these centres than was previously appreciated. Neurosecretory centres may thus also function as chemical brains wired by neuropeptide signalling where the specificity is derived not from direct synaptic connections but by peptide-receptor matching. Given a high enough number of specific peptide-receptor pairs and allowing the possibility of combinatorial signalling, it is theoretically possible to wire arbitrarily complex neural networks. This extends the concept of neurosecretory centres as organs that release peptides to influence external downstream targets.

An anatomically and molecularly distinct ‘chemical’ and ‘synaptic’ brain as we found in the *Platynereis* larva may have originated early in the evolution of the nervous system. This idea is consistent with the chimeric brain hypothesis, which posits the fusion of a neurosecretory and a synaptic nervous system early in animal evolution (Tosches and Arendt 2013). The neurosecretory part of the brain is directly sensory in *Platynereis*, possibly reflecting an ancestral condition for neurosecretory centres (Tessmar-Raible et al. 2007; Hartenstein 2006). The ANS also connects to the motor parts of the synaptic nervous system by specific projection neurons that are under peptidergic control. This connection may allow the translation of environmental cues that had been integrated by the chemical brain into locomotor output.

Neuropeptides are ideal molecules for chemical signalling due to their small size and high diversity. Many of the neuropeptides expressed in the *Platynereis* ANS and the vertebrate hypothalamus belong to ancient peptide families that evolved close to the origin of the nervous system (Jékely 2013; Mirabeau and Joly 2013). Neuropeptide genes have been identified in bilaterians, cnidarians, and even the early-branching metazoan *Trichoplax adhaerens,* that lacks a synaptic nervous system but has neurosecretory cells (Nikitin 2015; Smith et al. 2014). Chemical signalling by neuropeptides may have been important early during nervous system evolution in small metazoans where only peptide diffusion could coordinate physiological activities and behaviour. As metazoans grew larger, the coordination of their large complex bodies required control by a synaptic nervous system (Keijzer and Arnellos 2017). The separate origins of the chemical and the synaptic nervous system may still be reflected to varying degrees in contemporary brains. The *Platynereis* larval nervous system, and possibly the nervous system of other marine invertebrate larvae, shows a particularly clear segregation of non-synaptic and synaptic nervous systems.

The extent of non-synaptic signalling even in the relatively simple nervous system of the *Platynereis* larva highlights the importance of the combined study of connectomes and chemical signalling networks.

## Methods

### Electron microscopy reconstruction of *Platynereis* anterior neurosecretory plexus

We reconstructed the circuitry of cells in the anterior plexus from an existing 3-day-old *Platynereis* serial section transmission electron microscopy (ssTEM) dataset we generated previously (Randel et al. 2015). The dataset consists of 4845 layers of 40 nm thin sections. Preparation, imaging, montage, and alignment of the dataset is described (Randel et al. 2015). The cells were reconstructed, reviewed, 3D visualized, and the resulting synaptic network was analysed in Catmaid (Saalfeld et al. 2009; Schneider-Mizell et al. 2016).

### Neuropeptide expression atlas in 2-day-old *Platynereis*

DIG-labelled antisense RNA probes were synthesized from clones sourced from a *Platynereis* directionally cloned cDNA library in pCMV-Sport6 vector (Conzelmann et al. 2013). Or PCR amplified and cloned into the vectors pCR-BluntII-TOPO or pCRII-TOPO. Larvae were fixed in 4% paraformaldehyde (PFA) in 1 X PBS with 0.1% Tween-20 for 1 h at room temperature. RNA *in situ* hybridization using nitroblue tetrazolium (NBT)/5-bromo-4- chloro-3-indolyl phosphate (BCIP) staining combined with mouse anti-acetylated-tubulin staining, followed by imaging with a Zeiss LSM 780 NLO confocal system and Zeiss ZEN2011 Grey software on an AxioObserver inverted microscope, was performed as previously described (Asadulina et al. 2012), with the following modification: fluorescence (instead of reflection) from the RNA *in situ* hybridization signal was detected using excitation at 633 nm in combination with a Long Pass 757 filter. Animals were imaged with a Plan-Apochromat 40x/1.3 Oil DIC objective.

We projected thresholded average gene expression patterns of >5 individuals per gene onto a common 2-day-old whole-body nuclear reference templates generated from DAPI (Asadulina et al. 2012). Thresholding was performed either manually or following (Vergara et al. 2017). Gene expression atlases were set up in the visualization software Blender (https://www.blender.org/) as described (Asadulina et al. 2015).

### Deorphanization of receptor-peptide pairs

*Platynereis* GPCRs were identified in a reference transcriptome assembled from cDNA generated from 13 different life cycle stages (Conzelmann et al. 2013). GPCRs were cloned into pcDNA3.1(+) (Thermo Fisher Scientific, Waltham, USA) and deorphanization assays were carried out as previously described (Bauknecht and Jékely 2015).

### Analysis of single cell transcriptome data

Fastq files containing raw paired-end RNA-Seq data for 107 single cells from the 48 hour old *Platynereis* larval episphere (Achim et al. 2015) were downloaded from ArrayExpress, accession number E-MTAB-2865 (https://www.ebi.ac.uk/arrayexpress/experiments/E-MTAB-2865/). Only those samples annotated as ‘single cell’ and with a corresponding spatial mapping prediction by Achim et al. 2015 (Achim et al. 2015) were used in our analysis.

Fastq files with raw paired end read data were loaded into CLC Genomics Workbench v6.0.4 (CLC Bio). Data was filtered to remove Illumina adapter primer sequences, low quality sequence (Quality Limit 0.05) and short fragments (less than 30 base pairs). Filtered data were mapped to the assembled *Platynereis* reference transcriptome (including only sequences with a BLASTX hit e-value <1e-5 to the SwissProt database, plus all previously described *Platynereis* genes, including 99 proneuropeptides, a total of 52,631 transcripts. Mapping was carried out in CLC Genomics Workbench v6.0.4 using the RNA-Seq Analysis function, with the following mapping parameters: paired distance 100 – 800 base pairs; minimum length fraction 0.8, minimum similarity fraction 0.9, maximum number of mismatches 2. The total number of reads mapped to each gene, normalized by gene length (reads per kilobase million (RPKM)) in each sample was assembled into one spreadsheet using the ‘Experiment’ function, and this spreadsheet was exported as an Excel file. RPKM data for each gene and sample were converted in Excel to counts per million (cpm) by dividing RPKM by with the total number of mapped reads in each sample and multiplying by 10^6^ followed by conversion to a log base 10.

Single cell samples were sourced from populations consisting of dissociated cells of several individual larvae (Achim et al. 2015), therefore some of the RNA-Seq samples could represent sequencing of the same cell from different larvae. To determine a cut-off for transcriptome similarity to merge samples representing the same cell from different individuals, we calculated the all-against-all pairwise correlation coefficients. Plotting these correlation coefficients as a histogram showed a prominent peak of highly correlated samples. We interpret these as samples deriving from the re-isolation of the same cell from different larvae. We used a cut-off of 0.95 Pearson correlation as a cut-off, above which we merged samples as representing the same cell by using the mean normalized expression value for each gene.

Sample names were imported as nodes into software for graph visualization and manipulation, Gephi.0.8 beta (http://gephi.org). Nodes were manually placed in position in a Gephi map in an approximate 2D representation of the 3D spatial predictions of each cell generated by Achim et al. (Achim et al. 2015)based on a whole-mount *in situ* hybridization gene expression atlas of 72 genes (http://www.ebi.ac.uk/~jbpettit/map_viewer/?dataset=examples/coord_full.csv&cluster0=examples/resultsBio.csv). Samples with predicted bilateral symmetry were represented as two mirror image nodes in the map (left and right), while samples with predicted asymmetry were represented as single nodes. Node position coordinates for each sample were saved and exported as .gexf connectivity file for use in generating virtual gene expression patterns and peptide-receptor connectivity maps (Figure 3 – source data 3).

Virtual expression patterns for each gene were generated using a custom perl script that converted normalized log10 gene expression values into node colour intensity in the Gephi map. Connectivity files for each peptide-receptor pair were generated by preparation of connectivity data files as .csv files where the geometric mean of normalized log10 peptide expression from the ‘sending cell’ and corresponding GPCR expression from the ‘receiving cell’ was used as a proxy for connectivity strength. These connectivity data .csv files were imported into Gephi to generate .gexf connectivity maps, and random node positions were replaced with node position coordinates for each cell from the spatial position .gexf map described above. An ‘all-by-all’ connectivity map representing the potential cellular signalling generated by all known peptide-receptor pairs was generated by adding the connectivity data from each peptide-receptor pair into a single connectivity file.

Connectivity maps were analysed for degree of connectivity, graph density, modularity, number of connected components, and clustering coefficient in Gephi.0.8 beta. Nodes in connectivity maps were coloured by weighted in-degree (average number of incoming edges per node, adjusted for connectivity strength). Edge thickness was used to represent connectivity strength (geometric mean of peptide x receptor normalized gene expression). Authority was assigned to nodes using a Hyperlink-Induced Topic Search (HITS). Higher authority values indicate nodes that are linked to greater numbers of other nodes, or in the case of our peptidergic signalling maps, neurons with the capacity to receive signals from and send signals to the greatest number of other cells. Following export from gephi as .svg files, corresponding virtual neuropeptide expression for each connectivity map was used to colour signalling cells by overlaying in Adobe Illustrator CS6 V6.0.0 (Adobe Systems Inc.)

### Immunohistochemistry

Whole-mount triple immunostaining of 2 and 3 day old *Platynereis* larvae fixed with 4% paraformaldehyde were carried out using primary antibodies raised against RGWamide neuropeptide in rat (CRGWamide) and achatin neuropeptide in rabbit (CGFGD), plus a commercial antibody raised against acetylated tubulin in mouse (Sigma T7451). Double immunostaining was carried out with primary antibodies raised against MIP, RYamide or DH31 neuropeptide raised in rabbit and commercial acetylated tubulin antibody raised in mouse. The synthetic neuropeptides contained an N-terminal Cys that was used for coupling during purification. Antibodies were affinity purified from sera as previously described (Conzelmann and Jékely 2012). Immunostainings were carried out as previously described (Conzelmann and Jékely 2012).

### Calcium imaging experiments

Fertilized eggs were injected as previously described (Conzelmann et al. 2013) with capped and polA-tailed GCaMP6 RNA generated from a vector (pUC57-T7-RPP2-GCaMP6s) containing the GCaMP6 ORF fused to a 169 base pair 5' UTR from the *Platynereis* 60S acidic ribosomal protein P2. The injected individuals were kept at 18°C until 2-days-old in 6-well-plates (Nunc multidish no. 150239, Thermo Scientific). Calcium imaging was carried out on a Leica TCS SP8 upright confocal laser scanning microscope with a HC PL APO 40x/1.10 W Corr CS2 objective and LAS X software (Leica Microsystems) GCaMP6 signal was imaged using a 488-nm diode laser at 0.5 - 4% intensity with a HyD detector in counting mode. 2-day-old *Platynereis* larvae were immobilized for imaging by gently holding them between a glass microscope slide and a coverslip raised with 2 layers of tape as spacer. Larvae were mounted in 10 μl sterile seawater. For peptide treatment experiments, individual larvae were imaged for 2 minutes in the plane of the RGWamide interneurons and MC cell to assess the state of the larval nervous system prior to peptide treatment. For treatment, larvae were imaged for a further 5.5 min. 10 μl 50 μM synthetic D-achatin (G{dF}GD) (final concentration 25 μM) dissolved in seawater was added to the slide at the 1 minute mark by slowly dripping it into the slide-coverslip boundary, where it was sucked under the coverslip. As a negative control, larvae were treated with synthetic L-achatin (GFGD) at the same concentration. Receptor deorphanization experiments have shown that the D-form of achatin activates the achatin receptor GPCR, whereas the L-form of achatin does not (Bauknecht and Jékely 2015). D-achatin response was recorded from 12 larvae, and L-achatin response was recorded from 6 larvae. Calcium imaging movies were analysed with a custom Fiji macro and custom Python scripts. Correlation analyses were created using Fiji and a custom Python script (Verasztó et al. 2017).

## Acknowledgments

We thank Luis Bezares, Martin Gühmann and Nobuo Ueda for comments on the manuscript. The research leading to these results received funding from the European Research Council under the European Union’s Seventh Framework Programme (FP7/2007-2013)/ European Research Council Grant Agreement 260821. The research was supported by a grant from the DFG - Deutsche Forschungsgemeinschaft (Reference no. JE 777/1).

---

**Supplement 1 to Figure 1.**
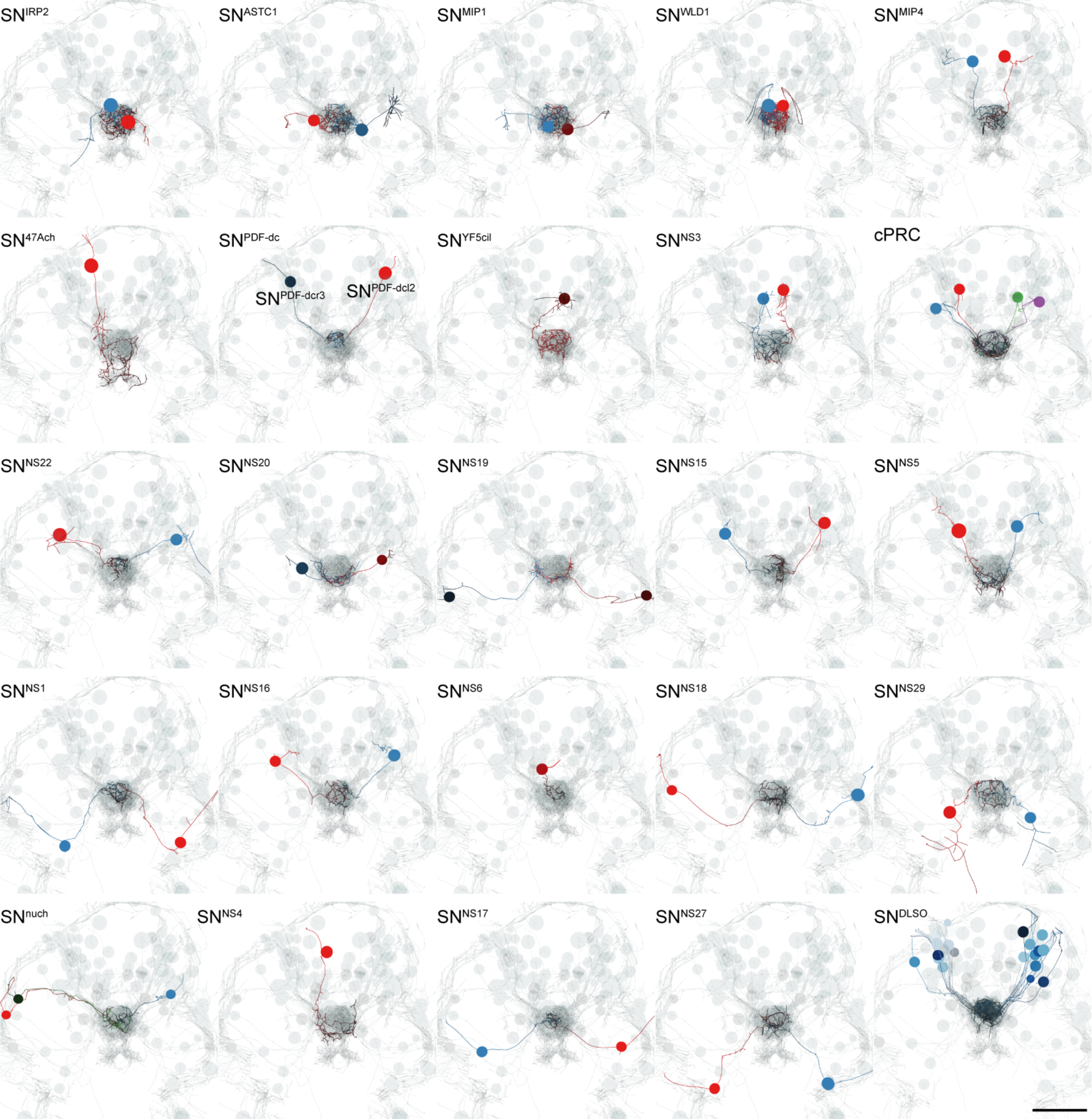
TEM reconstruction of ANS neurons. Bilaterally symmetric pairs and single asymmetric sensory neurons are shown in anterior view. Axons and dendrites appear as lines and cell body positions are represented by spheres. The reconstructed ciliary band cells and all ANS cells (grey) are included to show the perimeter of the head. Scale bar: 30 μm.

**Supplement 2 to Figure 1.**
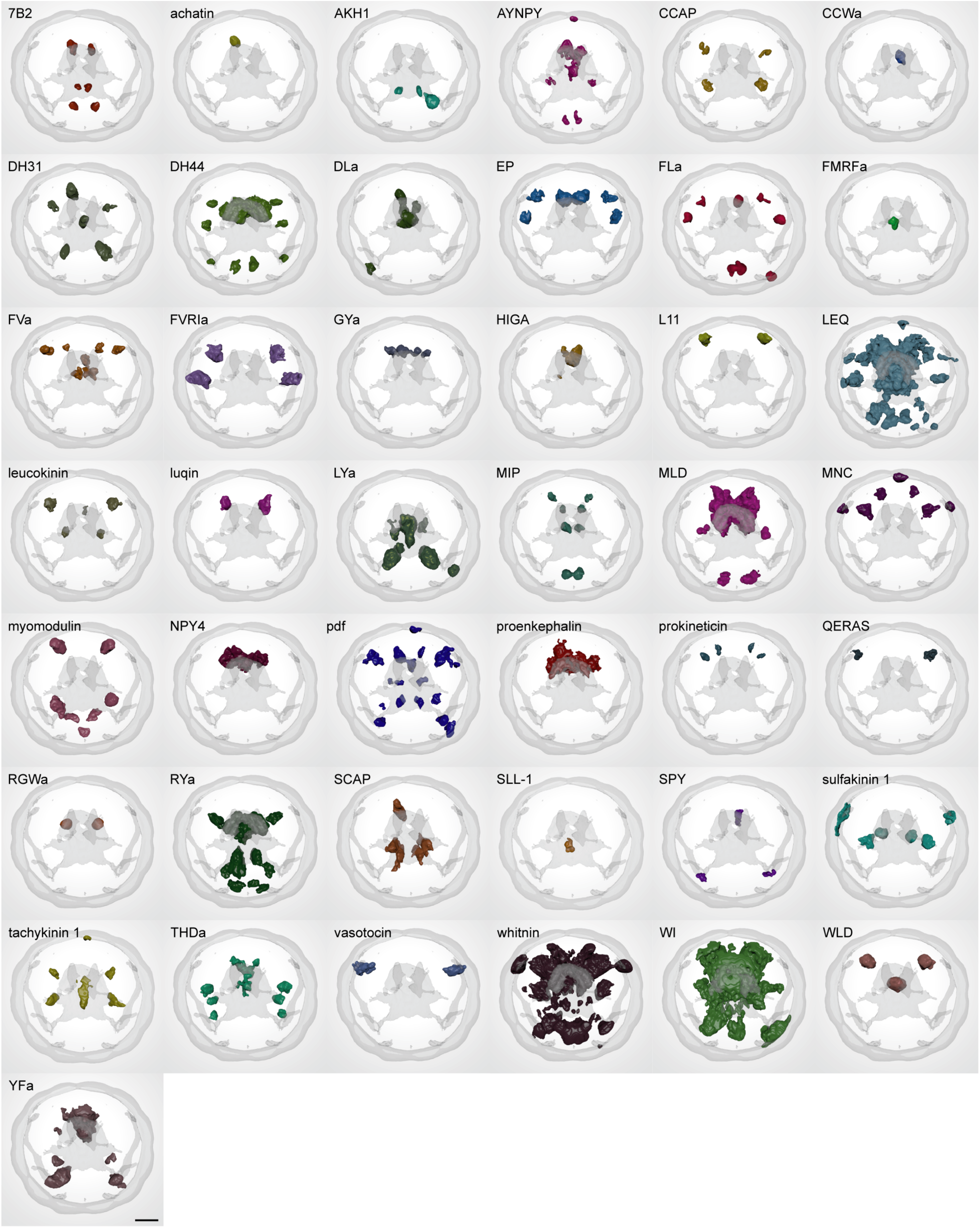
Average gene expression patterns of individual proneuropeptide from a whole-body gene expression atlas from 2-day-old larvae, anterior view. Scale bar: 30 μm.

**Supplement 3 to Figure 1.**
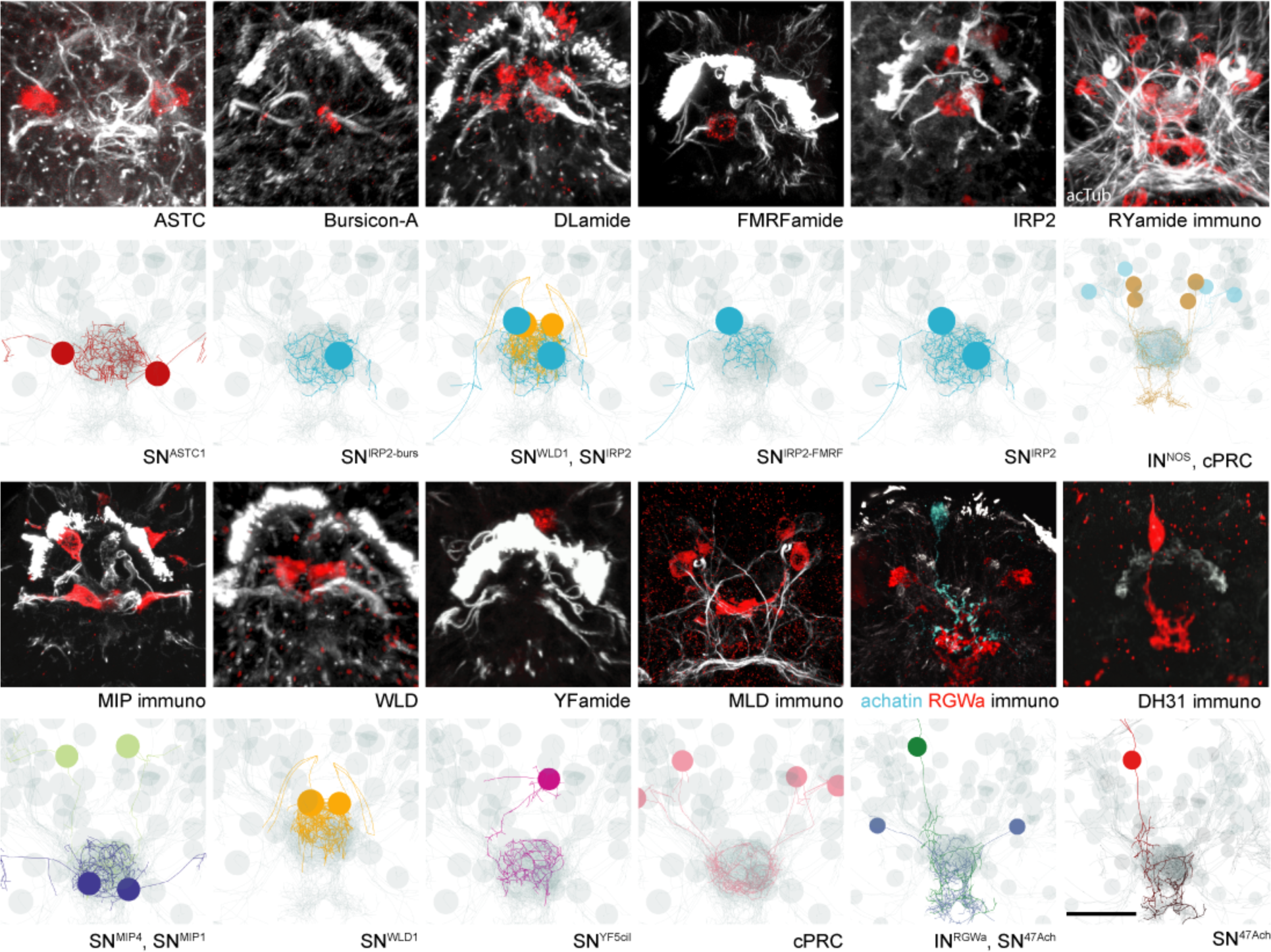
Expression of proneuropeptides in ciliated *Platynereis* larval sensory cells. Confocal microscope scans of whole-mount *in situ* hybridization of proneuropeptides (red) counterstained with anti-acetylated tubulin antibody (white). Close-up images show the apical sensory cilia of ANS neurons in 2-day-old larvae. The confocal stacks facilitated the assignation of neuropeptidergic identities to reconstructed neurons in the TEM connectome dataset. Scale bar: 25 μm.

**Supplement 1 to Figure 3.**
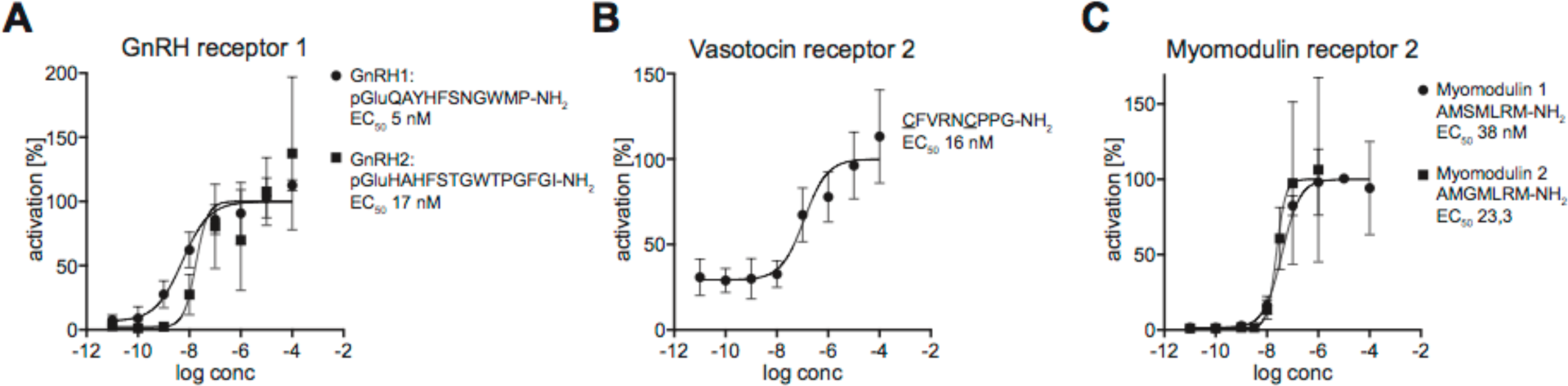
Dose-responsive curves of *Platynereis* deorphanized GPCRs treated with varying concentrations of peptides. Data, representing luminescence units relative to the maximum of the fitted dose-response curves, are shown as mean +/- SEM (n = 3). EC_50_ values and peptide sequences are shown beside each graph.

**Supplement 2 to Figure 3.**
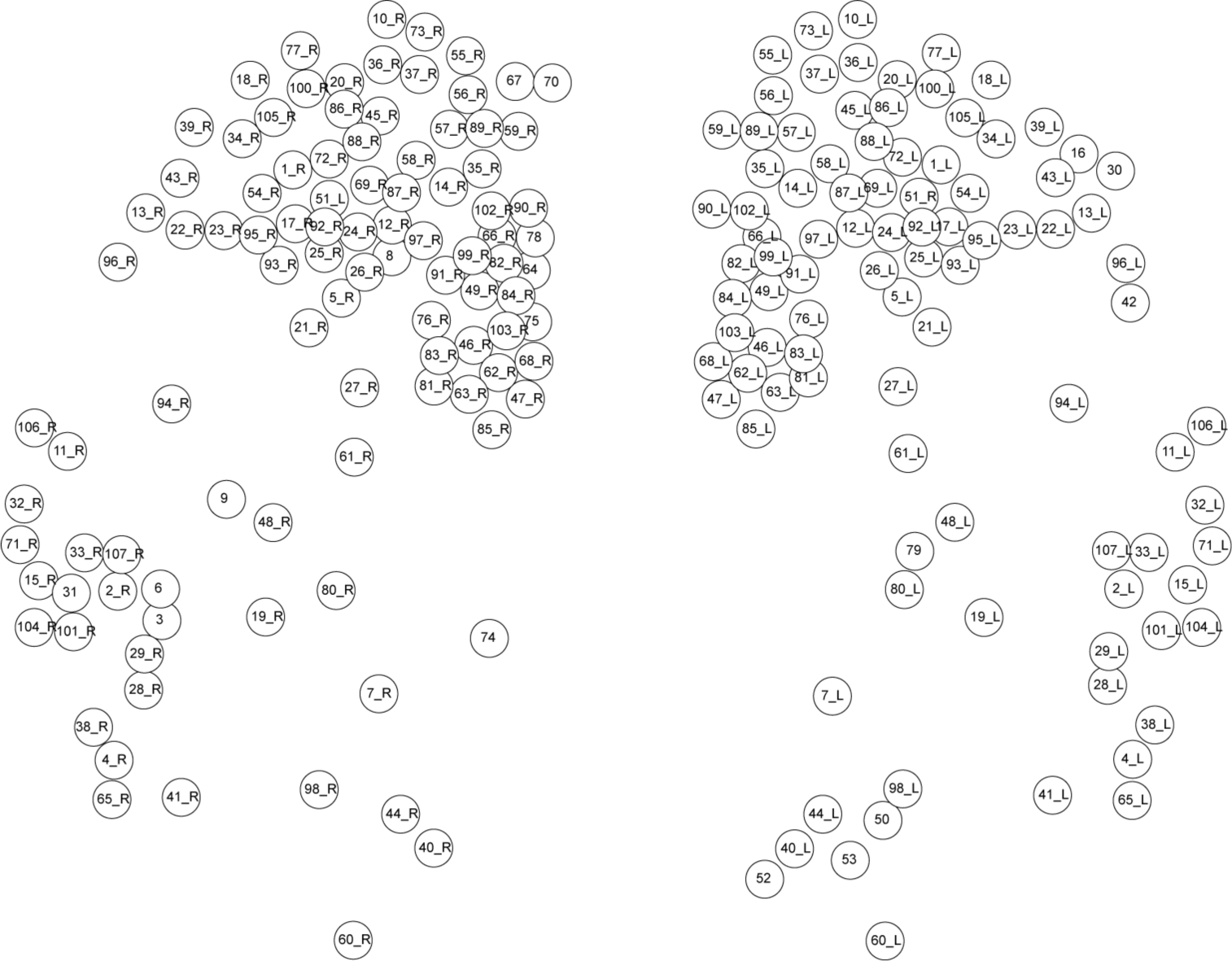
Spatial map of single cell RNA-Seq data from Achim et al. (Achim et al. 2015). The position of each node on the map is an approximation based on the spatial predictions of each RNA-seq sample generated by Achim et al. from comparisons of transcriptome expression with a whole mount *in situ* hybridization gene expression atlas of 72 genes. The correspondence of node IDs to original sample IDs from Achim *et al.* 2015 is listed in Supplementary table1.

**Supplement 3 to Figure 3.**
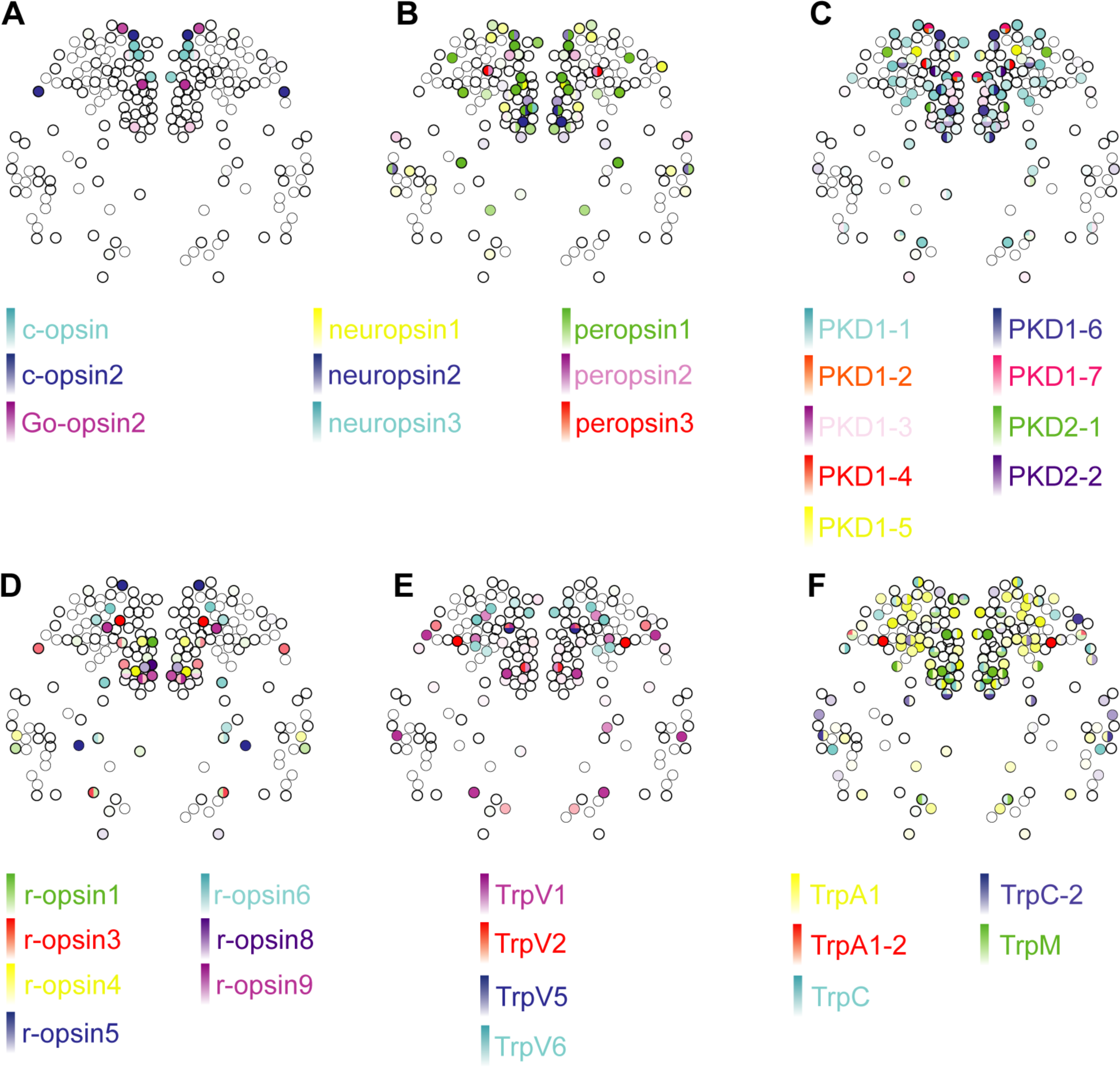
Expression of opsins and Trp channels in the *Platynereis* larval head. Colour intensity reflects relative normalized gene expression levels. Cells of the apical nervous system (i.e., cells expressing any combination of *Phc2, dimmed, Otp* and *nk2.1)* are indicated by a bold border.

**Supplement 4 to Figure 3.**
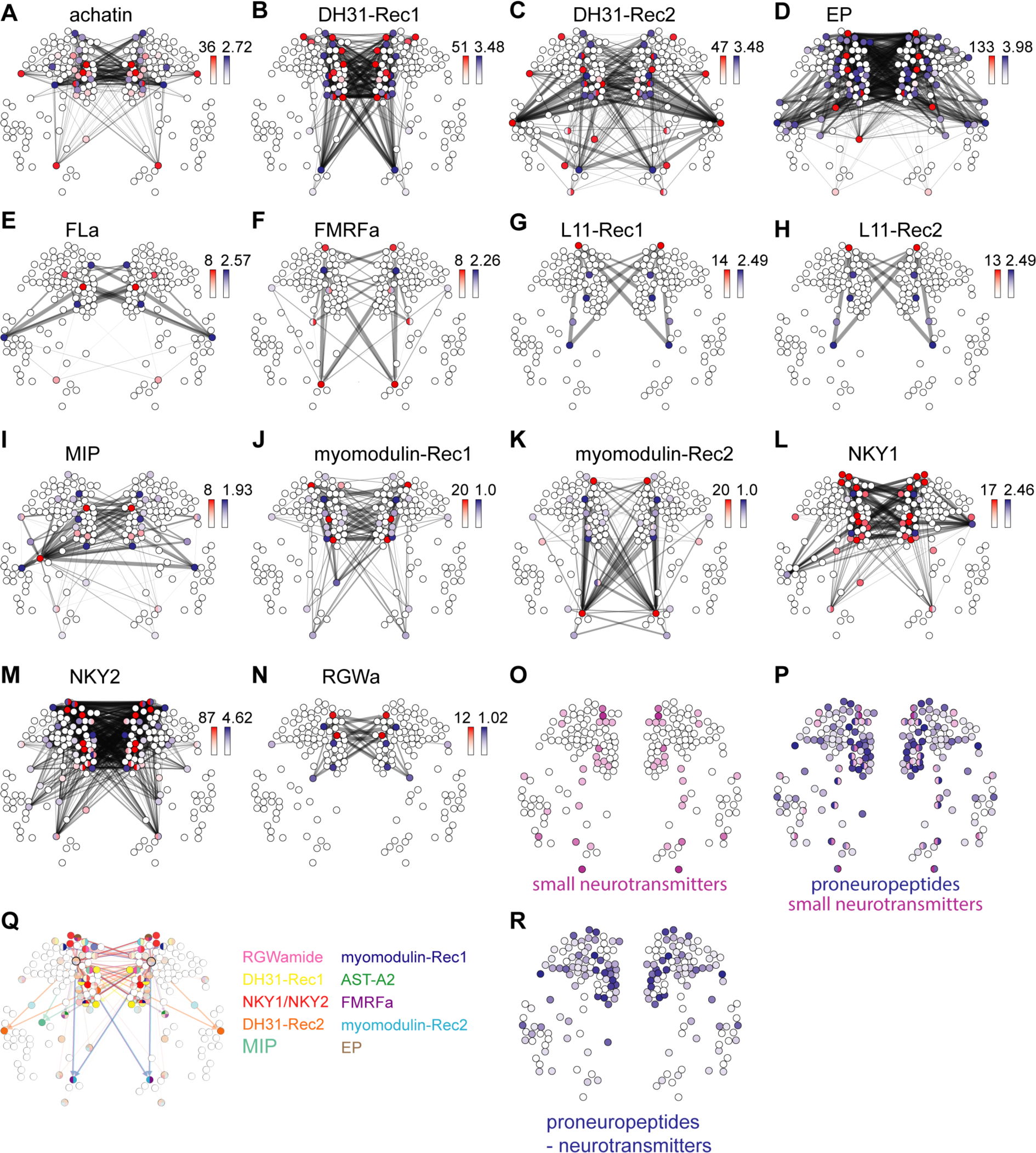
Neuropeptide-GPCR chemical connections in the *Platynereis* larval head. (**AN**) Connectivity maps of individual neuropeptide-GPCR pairs, coloured by weighted in-degree (red) and proneuropeptide log_10_ normalized expression (blue). Arrows indicate direction of signalling. Arrow thickness determined by geometric mean of log_10_ normalized proneuropeptide expression of signalling cell and log_10_ normalized GPCR expression of corresponding receiving cell. (**O**) Expression map of small neurotransmitter synthesis markers. (**P)** Expression map of proneuropeptides and small neurotransmitter markers. (**Q**) Multichannel signalling from a highly peptidergic cell. (**R**) Expression map of proneuropeptides shown for those cells that do not express small neurotransmitter markers.

**Supplement 1 to Figure 4.**
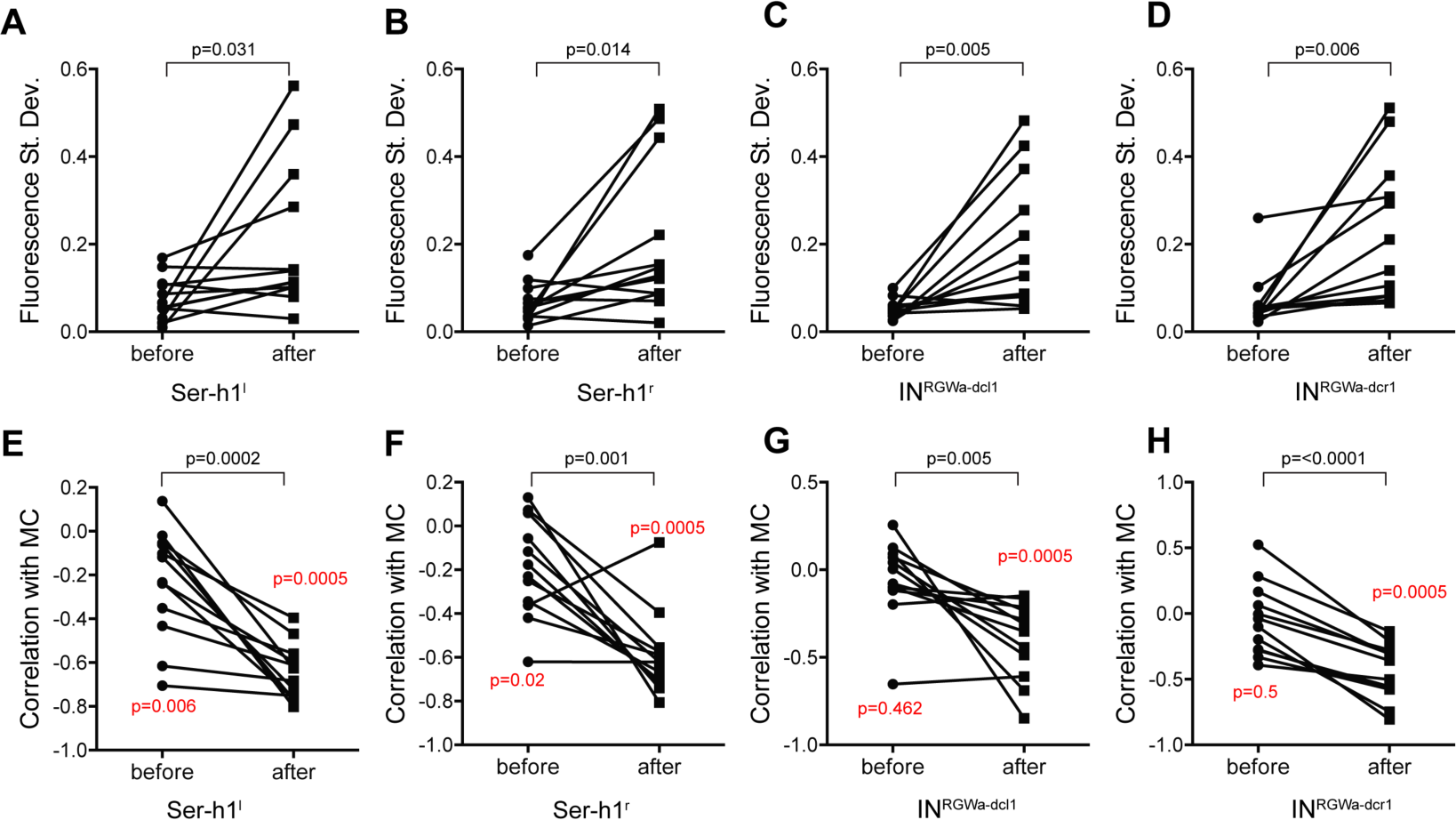
Standard deviation in fluorescent calcium signal of the (**A**) Ser-h1^1^, (**B**) Ser-h1^r^, (**C**) IN^RGWa-dcl1^ and (**D**) IN^RGWa-dcr1^ neurons in 2-day-old larvae before and after treatment with 25 μM D-achatin. (**E**-**H**) Correlation of neuronal calcium signals of (**E**) Ser-h1^l^, (**F**) Ser-h1^r^, (**G**) IN^RGWa-dcl1^ and (**H**) IN^RGWa-dcr1^ with calcium signals measured from the MC cell in 2-day-old larvae before and after treatment with 25 μM D-achatin. Individual data points represent different larvae before and after treatment. P-values of pairwise t-tests are shown above the square brackets. In (**E**-**H**) a Wilcoxon Signed Rank Test was also used to test if medians area significantly different from 0. The p-values are shown next to the data points.

## Supplementary Table 1

.xls file containing ANS synaptic connectivity spreadsheet, node ID to cell ID key, chemical network parameters, and log10 normalized expression values from mapping of single cell data for neuropeptides, GPCRs, sensory genes and neurotransmitter synthesis enzymes in separate worksheets.

## Video 1

EM reconstruction of the anterior nervous system in a 72-hr post fertilization *Platynereis* larva. The head ciliary band (grey) and the adult eye photoreceptor cells (transparent blue) are shown for reference.

## Author contributions

E.A.W and G.J. designed the study. E.A.W did single cell mapping, calcium imaging experiments, immunostaining and *in situ* hybridizations. C.V. and G.J. reconstructed neurons. S.J. did *in situ* hybridizations and generated the neuropeptide atlas. R.S. generated the EM dataset. P.B. deorphanized neuropeptide GPCRs. E.A.W. and G.J. wrote the paper.

